# Plastid degeneration in *Tillandsia* (Bromeliaceae) provides evidence about the origin of multilamellar bodies in plants

**DOI:** 10.1101/038158

**Authors:** Wouter G. van Doorn, Alessio Papini

**Affiliations:** Mann Laboratory, Department of Plant Sciences, University of California, Davis, California 95616, USA; Dipartimento di Biologia, Università di Firenze, Via La Pira 4, 50132 Florence, Italy; e-mail

**Keywords:** autophagy, chloroplast, multilamellar bodies, plastids, vacuole

## Abstract

Vesicle-like structures containing several to numerous concentric membranes, called multilamellar bodies (MLBs), are present both in animal and plant cells. The origin of MLBs in animal cells has been elucidated partially, while that of plant MLBs is unknown. MLBs in plant cells are present in the cytoplasm, at the interface of cytoplasm and vacuole, and inside vacuoles. This suggests that they become transported from the cytoplasm to the vacuole. The function of plant MLBs thus seems transfer of cellular membranes to the vacuole. Although it is often impossible to discern whether they have a single or a double outer membrane, in some examples a double outer membrane is present. This might suggest autophagic/mitochondrial/plastidial origin. Membrane structures similar to those in MLBs have not been described, apparently, in mitochondria. By contrast, structures similar to MLBs are found in autophagous structures and in degenerating chloroplasts and other plastids. The data might suggest the hypothesis that plant MLBs derive from autophagous structures and/or from plastids.

## Introduction

Multilamellar bodies (MLBs) have been shown in animal and in plant cells, and are also found in in protozoa (Paquet et al., 2013). In animals, MLBs contain several concentric membrane layers, and are bound by a single outer membrane. MLBs are found in many cell types, and are involved in storage of proteins and lipids. They also often function in secretion of lipids and other molecules to the cell surface (Schmitz and Müller, 1991; Hariri et al., 2000; Lajoie et al., 2005; Paquet et al., 2013). For example, in lung type II alveolar cells, MLBs are secreted to the exterior. This results in the deposition of lipidic surfactant molecules that regulate the surface tension at the air-lung interface (Lajoie et al., 2005). The hydrophobic film that is deposited on the stomach surface also originates from MLBs and so does the hydrophobic water-protective barrier of the skin (Schmitz and Müller, 1991; Raymond et al., 2008; Moreno et al., 2008).

Animal MLBs derive from the trans-Golgi network (Raymond et al., 2008). They also have been suggested to be formed by autophagy (Hariri et al., 2000; Lajoie et al. 2005) MLBs, have been interpreted to store, transport and secrete, and possibly digest, endosomal and/or autophagic cargo (Dermaut et al., 2005).

As in animals, plant MLBs consist of several concentric membranes, packed together in a vesicle-like compartment. Plants MLBs have been observed in the cytosol and in vacuoles, but have apparently not been reported to deposit their cargo in the apoplast. In contrast to MLBs in animals, no information seems available on the origin of the ones in plants. We here discuss a number of possibilities on the basis of observations in *Tillandsia* (Bromeliaceae) of the anther tapetum, a tissue undergoing degeneration processes in its last phase of development, eventually leading to programmed cell death (Papini et al. 1999). The frequent presence of MLBs in this tissues at different stages of development will be the basis to hypothesize a possible origin of plant MLBs.

## Materials and methods

### Plants

One of the authors collected *Tillandsia* spp. (mainly *Tillandsia albida* Mez et Purpus) in Mexico in 1991 and 1997 in Mexico D.F. and in the Hidalgo state. The plants were maintained as living specimens in the Botanical Garden of Florence (Italy) until nowadays. Flowers of *Tillandsia* were cut a different heights (and different developmental stages) of the inflorescence. Approximately ten flowers were examined at each stage of development. We cut also leaves sections for observing efficient chloroplasts.

We cut transverse sections (0.5 mm thickness, about three per flower) of the anthers and the same size of the leaf with a razor blade. These sections were fixed overnight in 1.25 % glutaraldehyde at 4° C in 0.1 M phosphate buffer (pH 6.8), then post-fixed in 1% OsO_4_ in the same buffer for 1 hr. After dehydration in an ethanol series and a propylene oxide step, the samples were embedded in Spurr’s epoxy resin (Spurr 1969). Transverse sections approximately 80 nm thick were cut with Reichert-Jung ULTRACUT E ultramicrotome equipped with a diamond knife. The sections on copper grids were stained with uranyl acetate (Gibbons and Grimstone 1960) and lead citrate (Reynolds 1963), and then observed with a Philips EM300 TEM at 80 kV.

## Results

In the tapetum of Tillandsia at the tetrads stage of pollen development some MLBs could be found free in the cytosol, often in contact with small vacuoles (Fig. 1). During a later stage the cytoplasm of the tapetal cells tended to become more electron dense (Fig.s 2 and 3). In this stage many MLBs were observed entering a small vacuole (Fig. 2) or already included in it (Fig. 3). In this latter situation the MLB appeared to have a central area with a lower electron density but always containing membranes. Other membranes not organized as MLBs were observed elsewhere in the vacuole. Vacuoles tended in some cases to fuse together to form larger structures containing MLBs (Fig. 4). MLBS proceding from the situation in Fig. 2 to Fig. 3 and 4 appeared to reduce their size. In some cases portion of cytoplasm and MLBs could be individuated contemporaneously within a medium sized vacuole (Fig. 5).

**Fig. 1.**
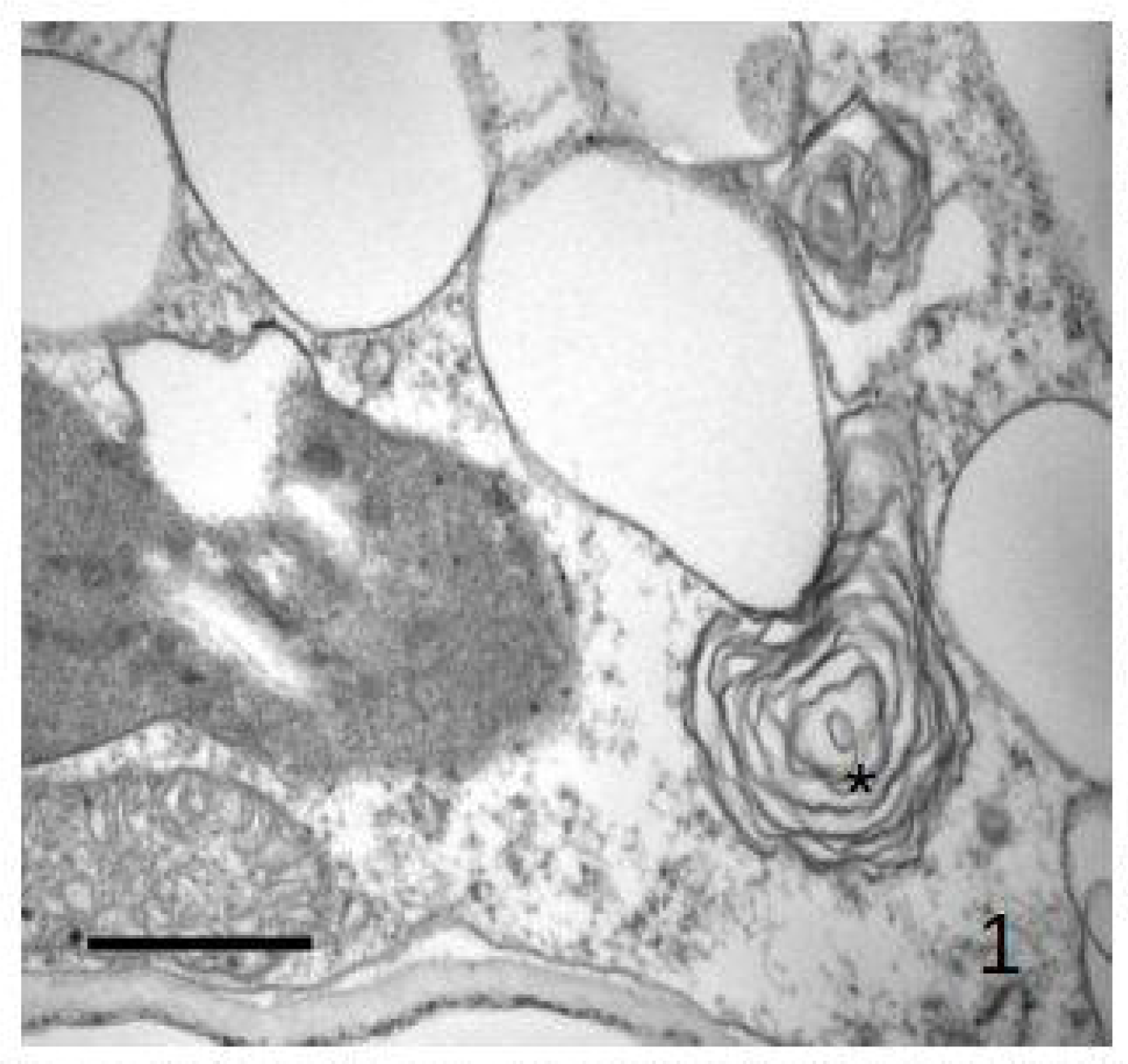
Multilamellar bodies in the cytosol (asterisk) in *Tillandsia albida* tapetum cell. Bar = 500 nm.

**Fig. 2.**
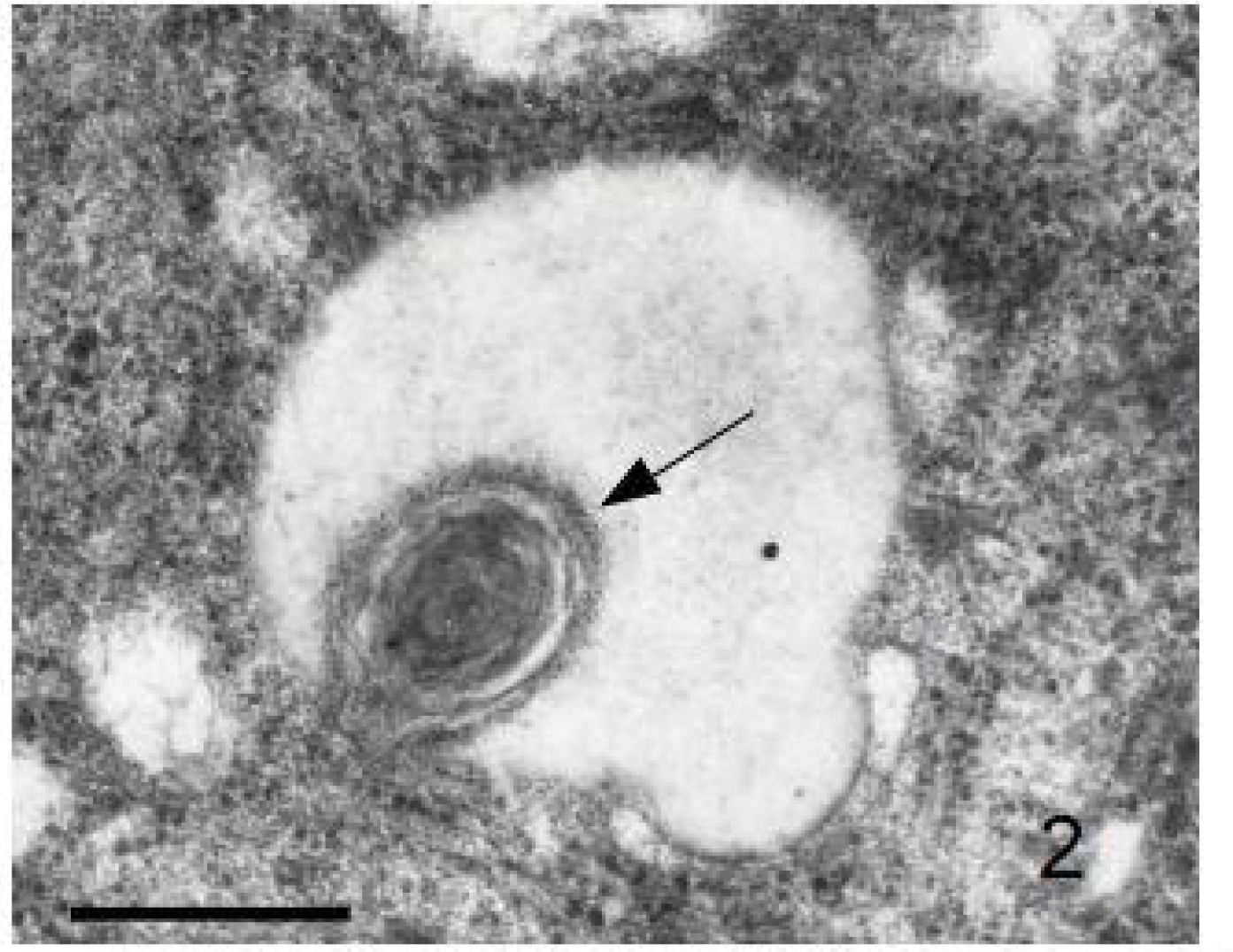
Multilamellar bodies entering the vacuole in *Tillandsia albida* tapetum cell. Bar = 1 μm.

**Fig. 3.**
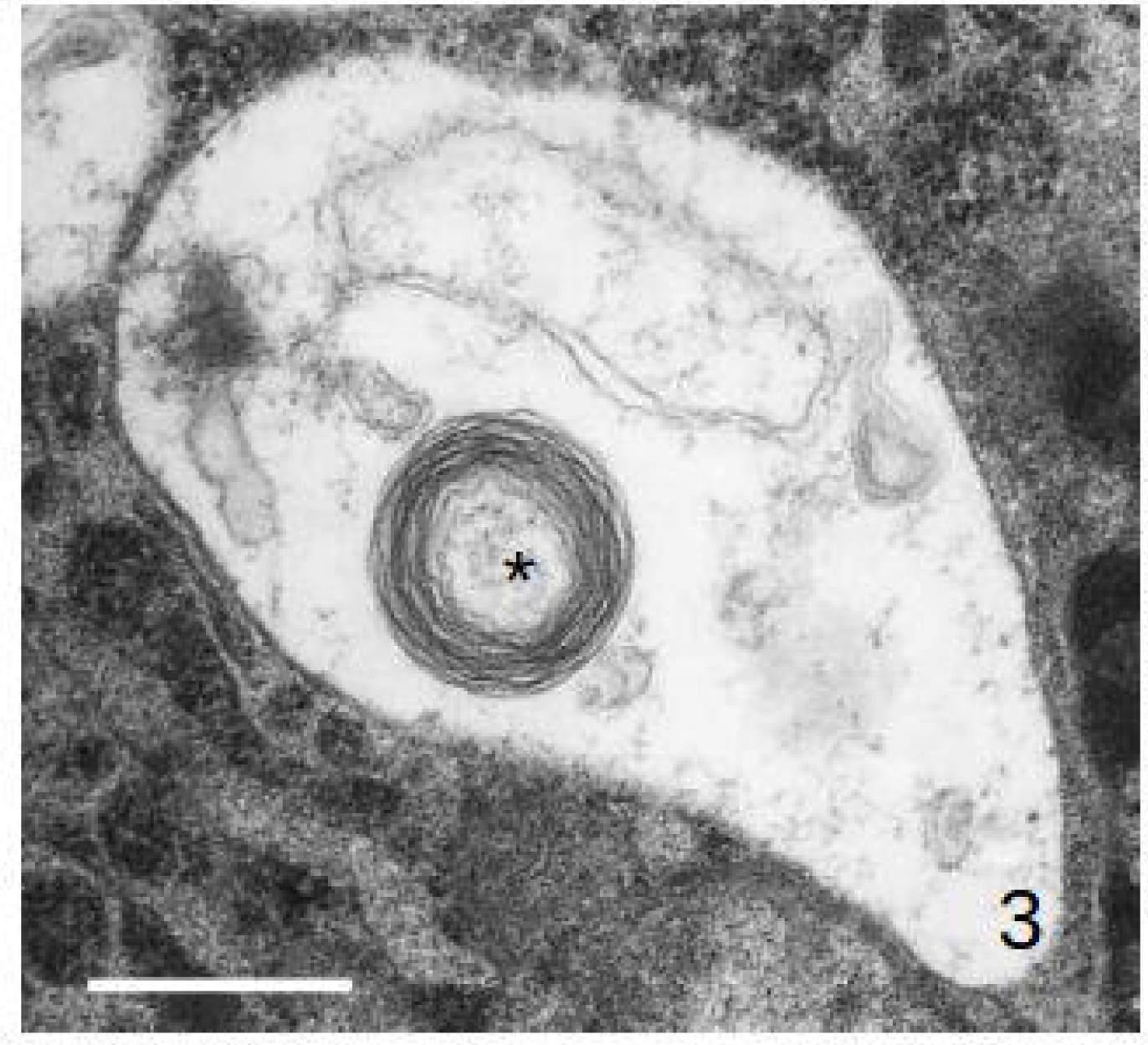
A single MLB in a vacuole Bar = 500 nm.

**Fig. 4.**
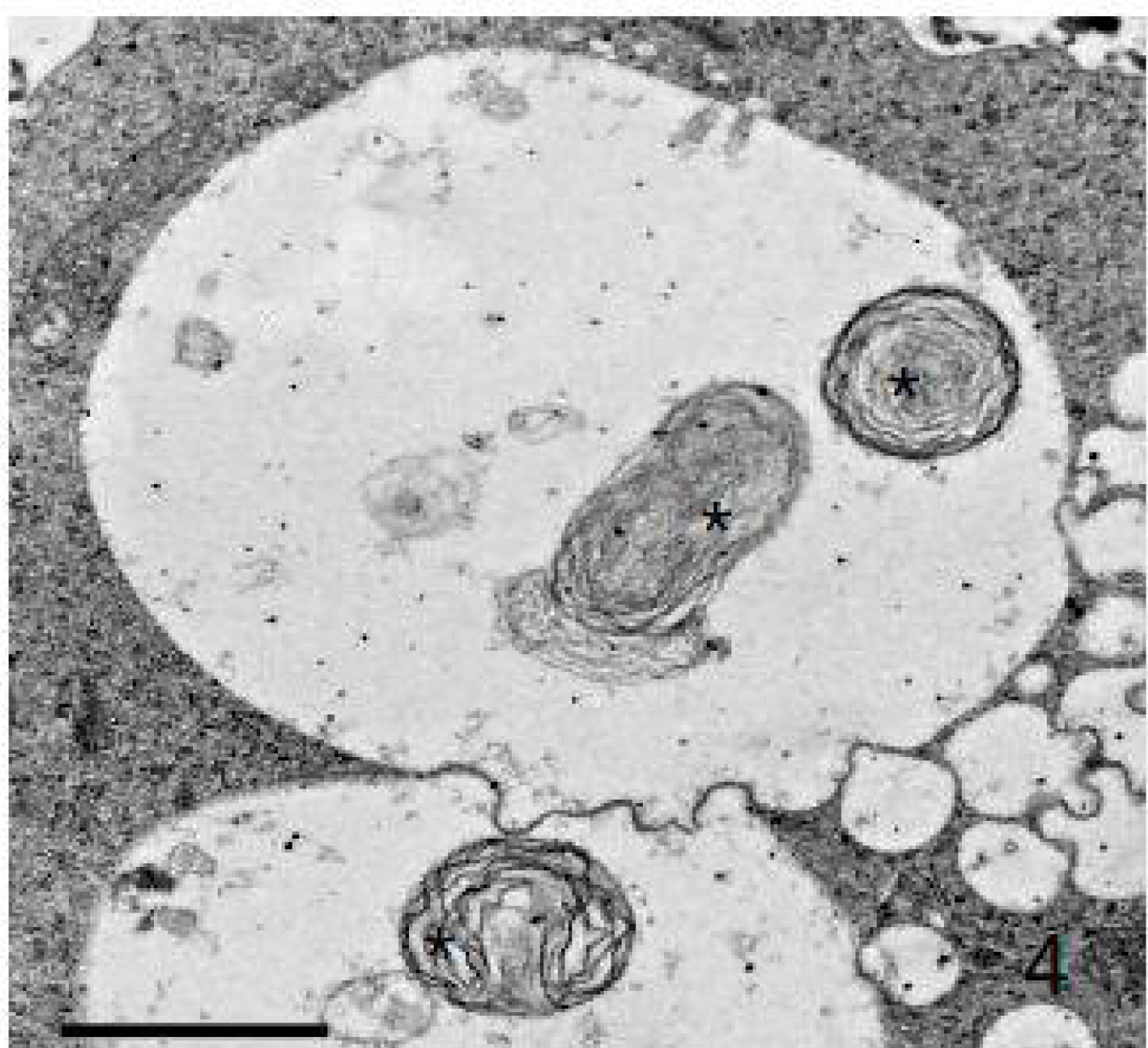
Vacuoles containing MLBs merging together Bar = 500 nm.

**Fig. 5.**
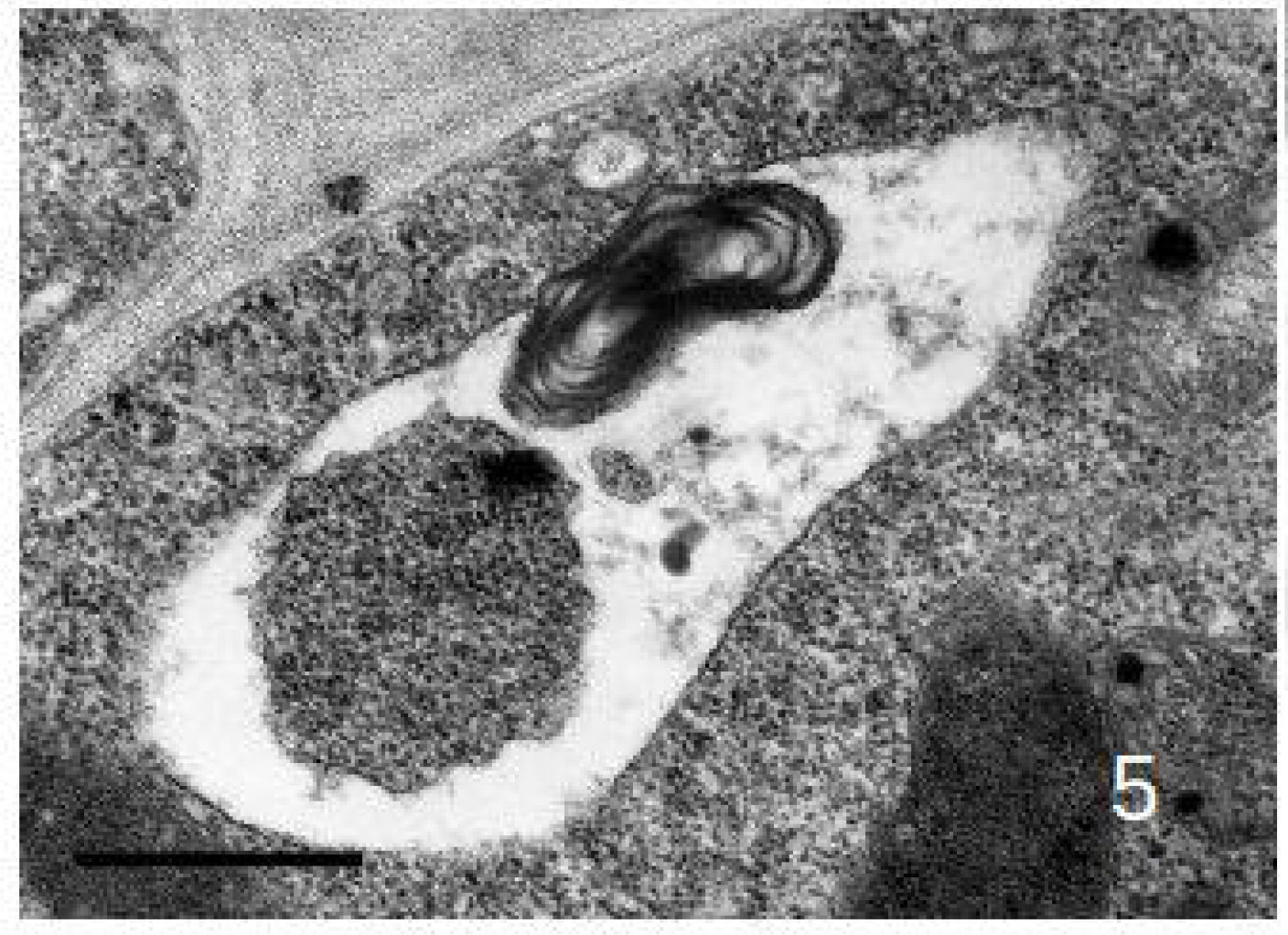
A portion of the cytoplasm and an MLB in a plant autophagosome/autolysosome in a *Tillandsia albida* tapetum cell. Bar = 500 nm.

In *T. albida* leaves, the guard cells of stomata showed normal chloroplasts containing starch and apparently normal thylakoids and some chloroplasts still containing starch but starting to dismantle their membrane system (Fig. 6).

**Fig. 6.**
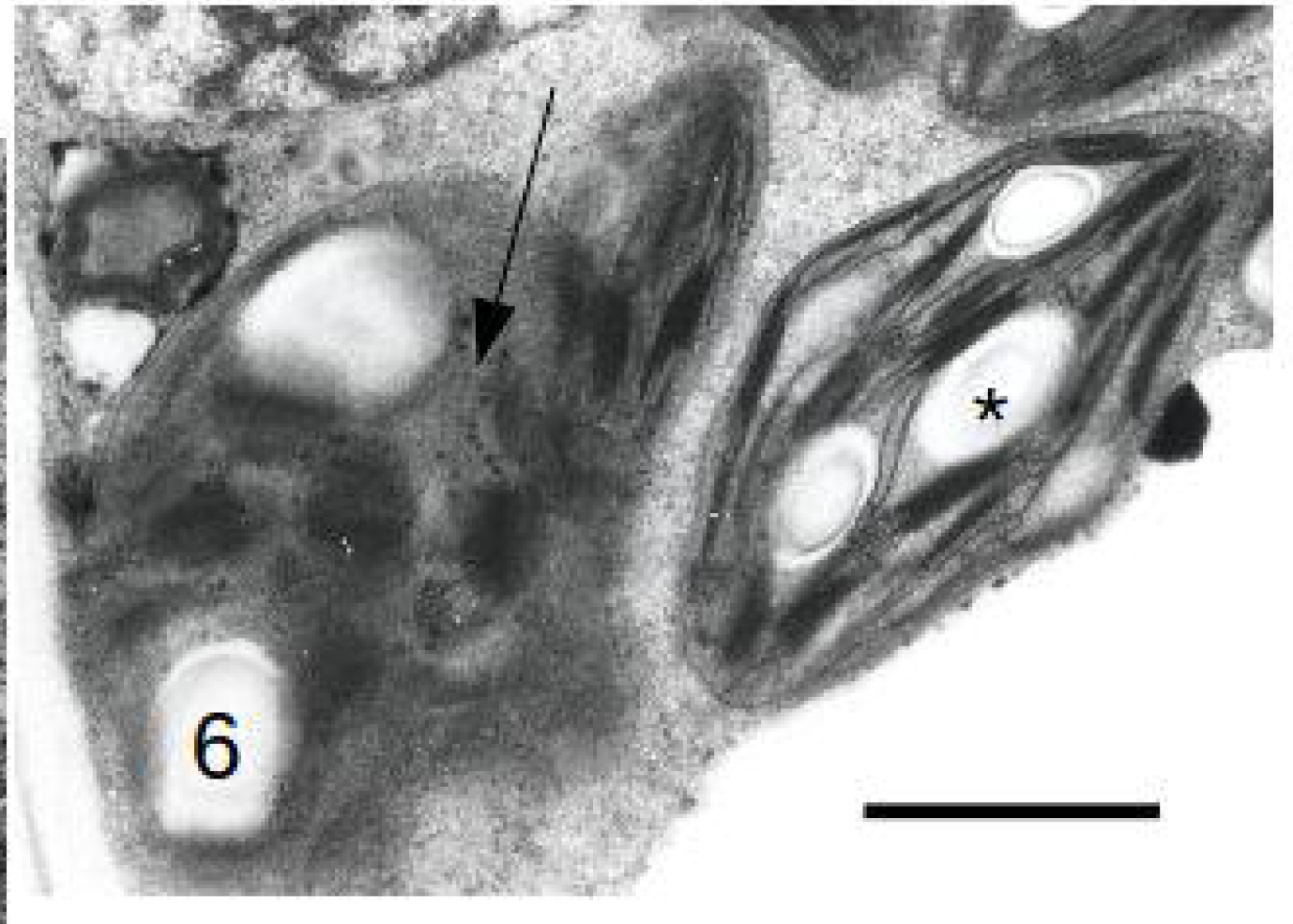
*T. albida* stomatal guard cell. Normal chloroplast (asterisk) early stages of a chloroplast degeneration (arrow). Bar = 200 nm.

At a later stage of tapetal development concentric membranes in plastids became frequent (Fig. 7). In some rare cases some plastoglobules appeared surrounded by a membrane.

**Fig. 7.**
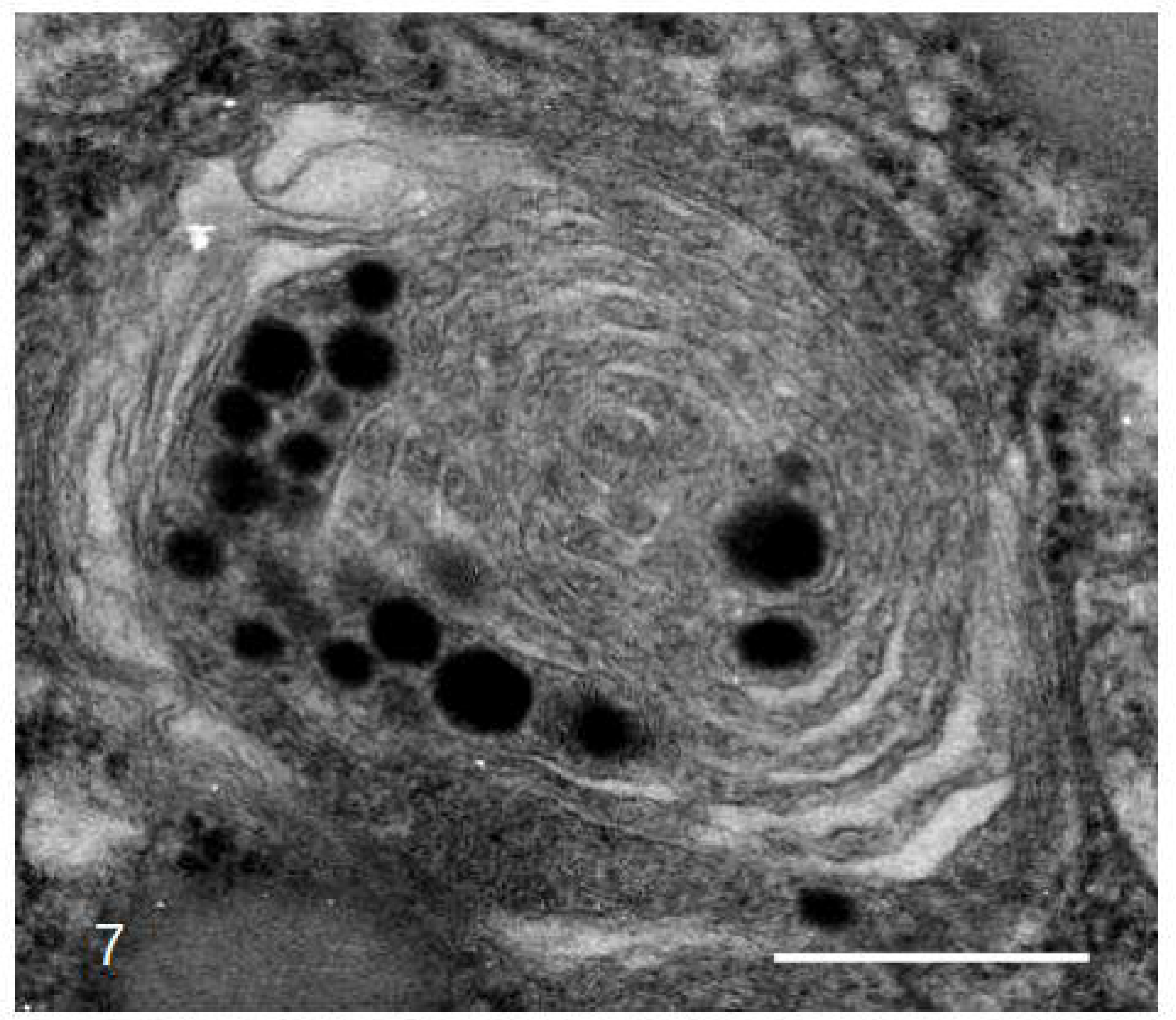
Concentric membranes in plastids of *T. albida* tapetum cells. Bar = 200 nm.

**Fig. 8.**
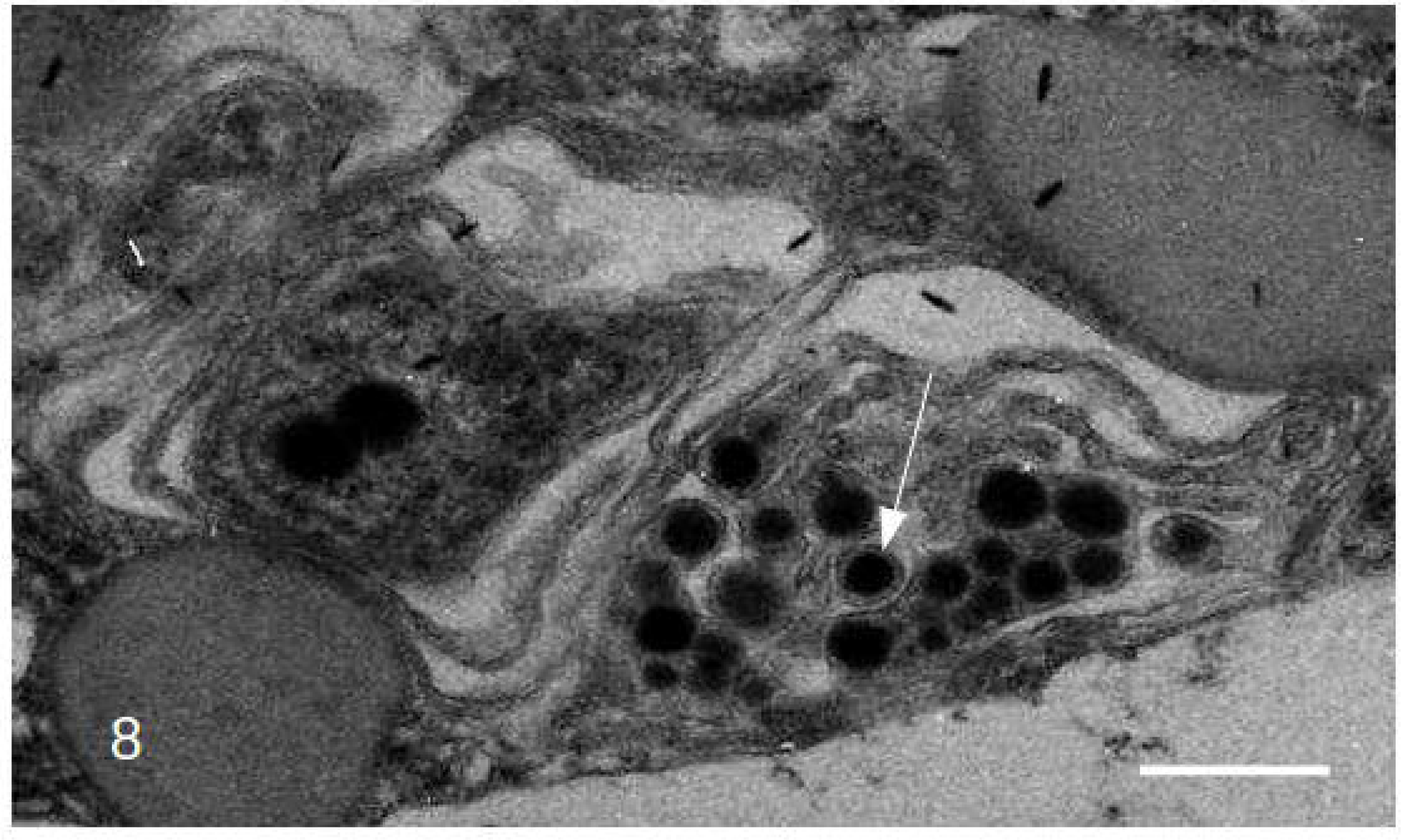
As in Fig. 7, but Some plastoglobules (arrow) appear surrounded by a membrane. Bar = 200 nm.

Some cells contained large vacuoles that engulfed large portions of protoplast including apparently organells with a continuous double membrane. Inside the organelle are also membranes (Fig. 9).

**Fig. 9.**
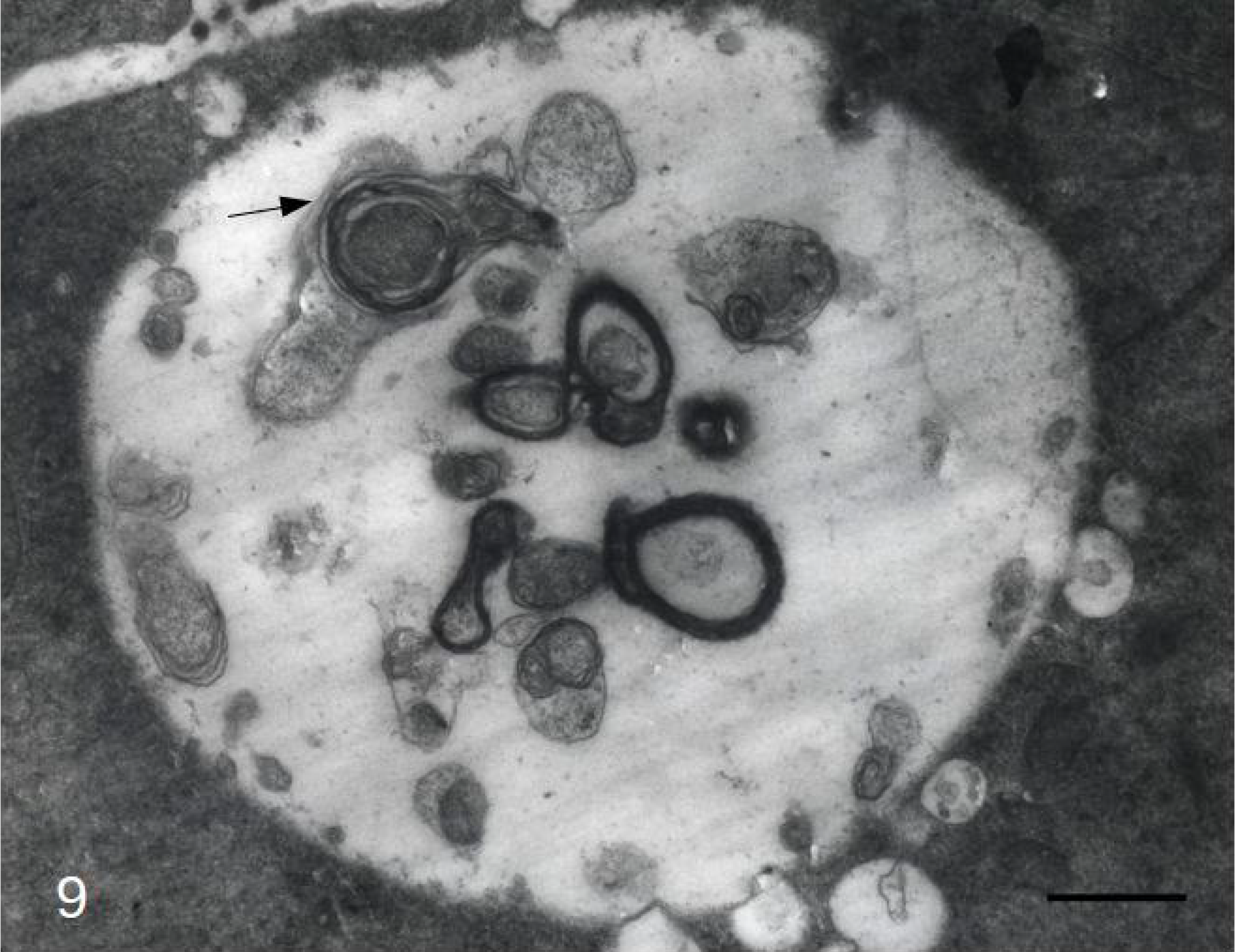
*T. albida* tapetum cells. Double membranes of MLBs and autophagous sequestration of plastids or MLBs. Autophagosome/autolysosome. The engulfed portions of the protoplasm contain what seem organelles, possibly plastids, with a continuous double membrane (arrow). Inside the organelle are also membranes. Bar = 2 μm.

Some plastids showed membrane-like structures possibly related to plastoglobuli and membranous structure with some electron-dense areas (Fig. 10). In other plastids concentric very uniform membranes accumulated (Fig. 11).

**Fig. 10.**
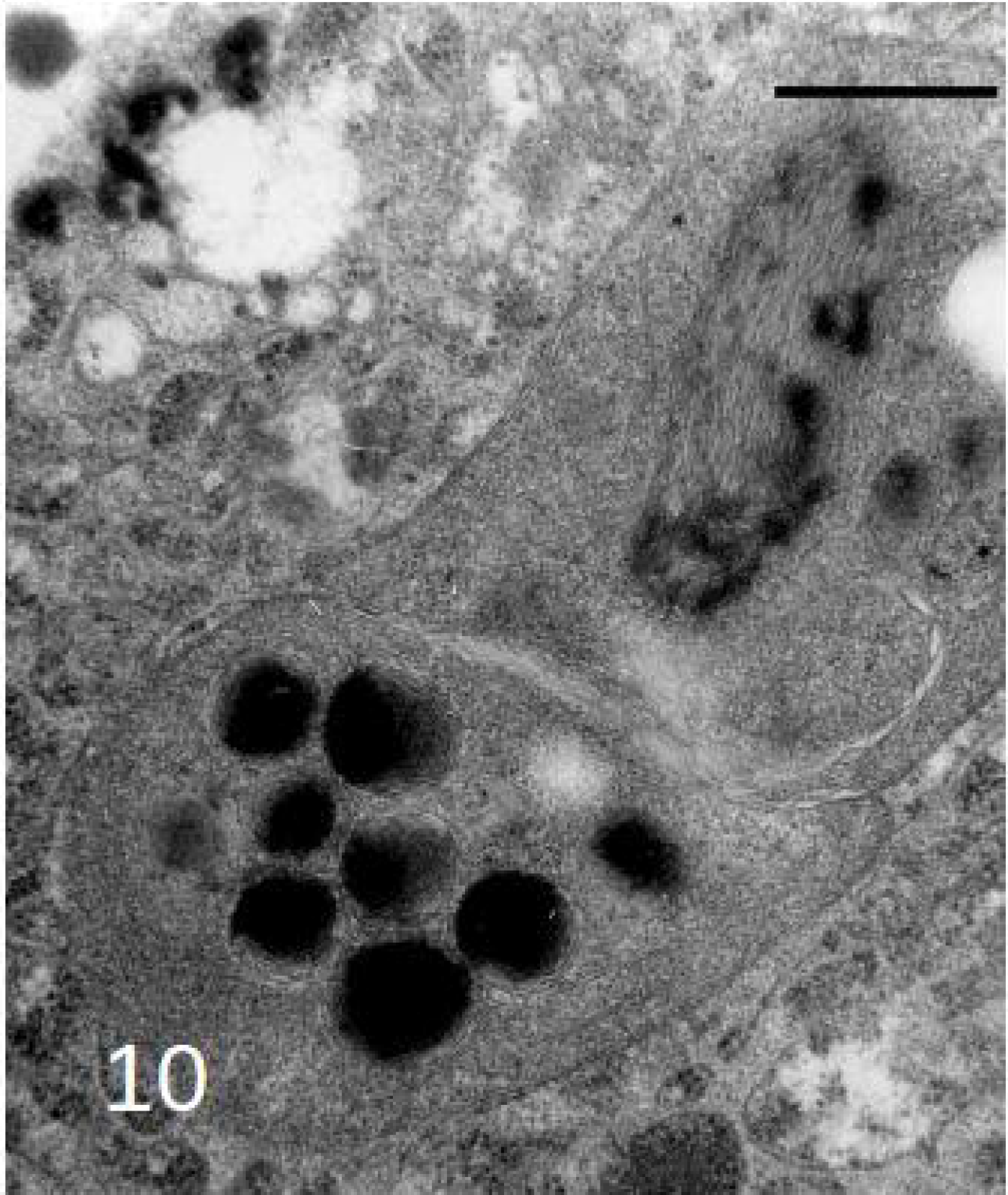
Concentric membranes in plastids of *Tillandsia albida* tapetum cells. Membrane-like structures possibly related to plastoglobuli (lower part of the micrograph) and membranous structure with some electron-dense areas (higher part of the micrograph). Bar = 500 nm.

**Fig. 11.**
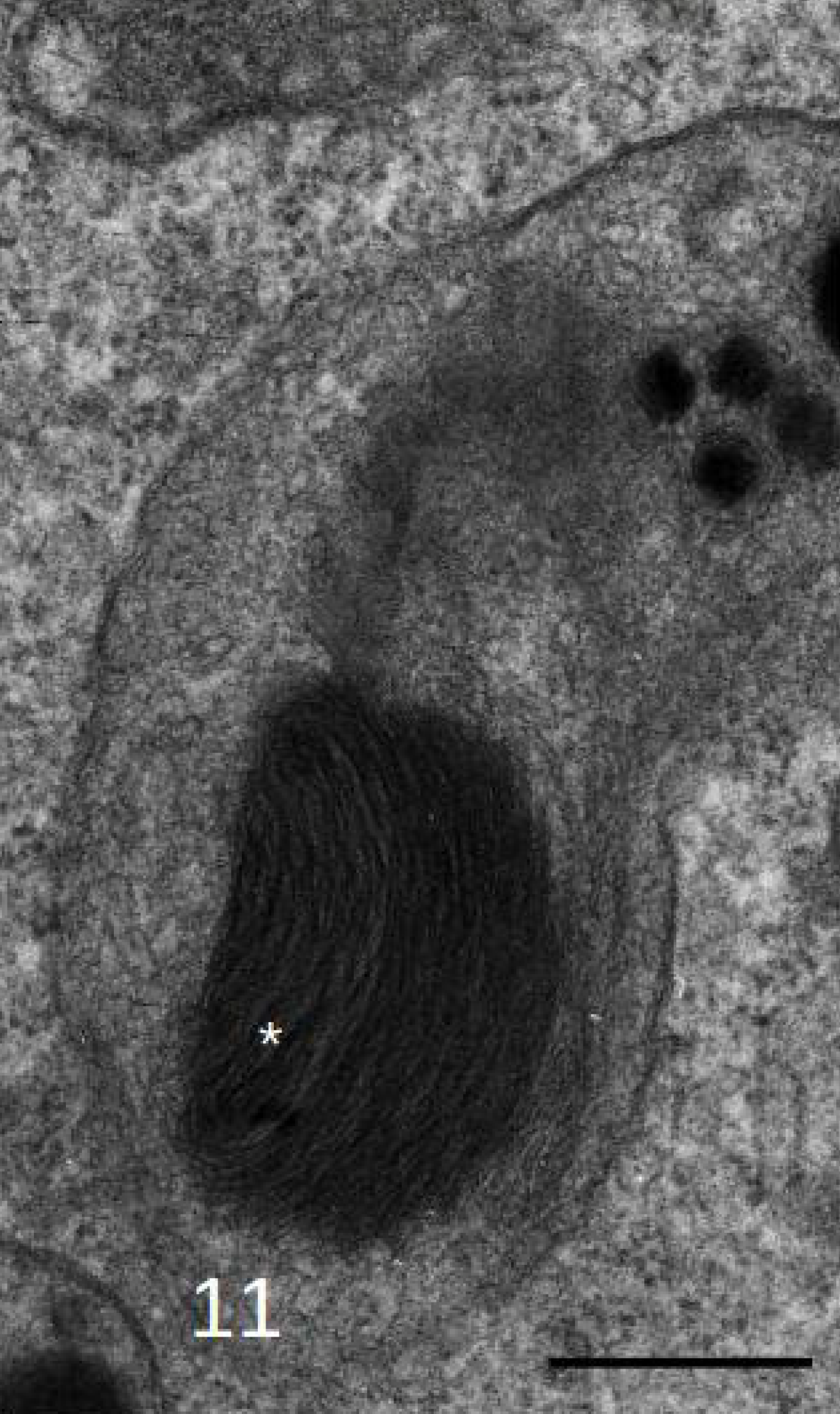
Concentric membranes in plastids of *Tillandsia albida* tapetum cells. Accumulated membranes that are rather uniform electron-dense form in a plastid (asterisk). Bar = 500 nm.

## Discussion

### Form, size and internal structure of plant MLBs; cellular localisation

Early authors called plant MLBs ‘myelin bodies’. Myelin refers to cells around neuronal axes, containing concentric membrane circles. Plants do not contain myelin (Adami and Aschoff, 1906; Nave, 2010).

Plant MLBs in plants vary in form, size and structure. They are usually round but can also be ovoid and oblong. In our work on tapetal cells in *Tillandsia albida* the largest diameter varied from 100 to 600 nm (Papini et al., 1999), but diameters up to 2400 nm have been reported (Schmitz and Müller, 1991; Fernández et al., 2013). The internal structure of plant MLBs varies in two ways: a) the number of stacked membranes and b) the membrane arrangement.

Membranes can be single or double. Examples of single membranes are found in plastids shown in Fig. 1. Double membranes were described in *Sarcocapnos pulcherrima* pollen grains (Fernández al. 2013). The origin of double membranes is not known. They might derive, for example, from degradative stages of mitochondria and/or plastids.

The MLBs shown in Fig. 1 are localised in the cytosol. Other MLBs have been found at the interface of cytosol and vacuoles, whereby they apparently protrude into the vacuole (Fig. 2). In still other examples the MLBs are observed inside vacuoles (Fig. 3,4). This sequence might suggest that MLBs in the cytosol can be transferred to vacuoles, where they become degraded. Plant MLBs also have been shown to become engulfed by autophagous structures (see below), which likely result in their degradation inside this structure.

### Are MLBs formed by autophagy?

Plant macroautophagy is different from that in animals. In animal cells a portion of the cytoplasm becomes surrounded by a double-membrane-bound structure that does not contain hydrolases. After sequestering a portion of the cytoplasm this organelle merges with a lysosome, which delivers hydrolases involved in breakdown of the portion of the cytoplasm. Thus in animals one phase of autophagy does not yet entail degradation. In plants the macroautophagous organelle contains hydrolases before sequestering a portion of the cytoplasm. Fig. 9 depicts an example of a plant autophagosome/autolysosome that sequesters portions of the cytosol, apparently containing organelles. The engulfed organelle has a double membrane, so might be plastidic or mitochondrial in origin. Inside this organelle are some more (double) membranes.

Similar structures were found by Rose et al. (2006) in isolated starving *Arabidopis thaliana* cell. Such an autophagic process might lead to a structure containing several concentric membranes, thus a MLB. However, it is not known how long these membranes might persist in such an autolytic vacuole. Plant macroautophagy tends to results in rather rapid breakdown of the sequestered material (van Doorn and Papini, 2013). If this is true, it would mean that plant autophagy will likely not add long-lived membranes to MLBs. If so, this would be an argument against an autophagous origin of MLBs. It cannot be excluded, however, that some membranes are hard to degrade and thus persist, hence that MLBs can - in principle - derive from macroautophagy, for instance in cells subjected to programmed cell death, as MLBs appearing in the scutellum of germinated wheat grains (Dominguez et al., 2012).

Whatever their origin (autophagic, plastidial, mitochondrial, or other), mature MLBs might be sequestered by macroautophagy (van Doorn and Papini, 2013). A possible example is shown in Fig. 5, where a macroautophagous structure has sequestered a portion of the cytoplasm and a MLB as well.

### MLBs derive from plastids?

Several authors noted a coincidence between chloroplast degradation during leaf senescence and the presence of multilamellar bodies in the cytoplasm or vacuole (Hurkman and Kennedy, 1975). Fig. 6 shows an example of a chloroplast. The internal membranes are arranged in thylakoids and grana. A thylakoid consists of two membranes closely positioned together, similar to the structure of the ER. Grana are stacks of thylakoids. At an early stage of disintegration the internal membranes structure changes, while plastoglobuli, electron dense bodies mainly containing lipids and proteins, appear or increase in number, as shown also by Evans et al. (2010), Biswal et al. (2013) and Sakuraba et al. (2014). The membranes in such plastids may show one major membrane swirl (Dennis et al. 1967) or several such swirls (Evans et al. 2010). In other examples of late stage chloroplasts only few membranes are left (Butler and Simon, 1971; Hurkman, 1979; Inada et al., 1998). During late phases of internal chloroplast degradation the organelle becomes considerably smaller (Ljubešic, 1968).

TEM micrographs were interpreted to show that whole chloroplasts, at some stage of internal breakdown, can be transported to vacuoles (Wrischer, 1973; Wittenbach et al., 1982; Minamikawa et al., 2001). However, these data are not convincing, as the location of the vacuolar membrane was not demonstrated. It therefore remains to be unequivocally demonstrated that whole chloroplasts can be transported to vacuoles (van Doorn and Papini, 2013).

The ultrastructure of chloroplasts and other plastids might suggest that they become MLBs. This appears, for example, in *Tillandsia* tapetum cells. The plastid in Fig. 7 shows several plastoglobuli and a concentric arrangement of membranes. These double membranes, which are similar to thylakoids, enclose relatively electron-dense material. In between the double membranes the plastid stroma is more electron-translucent. The plastid at the lower side of Fig. 8 shows plastoglobuli and a circular membrane structure similar to the one in Fig. 7. Above this plastid is another with only two plastoglobuli in the plane of section. This plastid has a few circular double membranes.Circular or bent membranes structures in plastids were found also in genera other than *Tillandsia.* As for plastids containing plastoglobuli of various size in a senescing *Nicotiana rustica* leaf (Ljubešic, 1968). Some circular membranes were also found in chloroplasts of senescing *Arabidopsis thaliana* leaves (Evans et al., 2010). Stacks of apparently rolled-up thylakoids have also been observed in chromoplasts of *Narcissus pseudonarcissus* flowers. These stacks consisted of about 5-10 double membranes, situated at the periphery of the organelle (Al-Babili et al., 1999). In chromoplasts of *Capsicum annuum* fruit single circular thylakoid sheets were observed as well as configurations of sheets which were either linear or rolled-up (Spurr and Harris, 1968, their Figs. 12-15; Laborde and Spurr, 1973, their Figs. 2, 7, 8, 12, 13).

Fig. 10 and 11 exhibit plastids with what are apparently plastoglobuli from which the electron-dense material (lipid) has been removed partially. Membrane-like structures have become visible. Similar membranous structures were found by Evans et al. (2010) in *Arabidopsis* mesophyll. Plastoglobuli have been associated not with membranes but with a single sheet of phospholipids, a half-membrane (Bréhélin et al., 2007; Lundquist et al., 2013). It is not clear if half membranes might be part of the typical rolled-up structures in MLBs. Lichtenthaler and Peveling (1967) showed membranous material in two organelles, most probably plastids as they also contain plastoglobuli. Such membraneous material might consist of half-membranes.

### Do MLBs derive from mitochondria?

The fate of plant mitochondria is still an enigma. A previous investigation in *Dendrobium* suggested that the internal mitochondrial membranes disappear during late stages of development (Kirasak et al., 2010). These data do not exclude that plant mitochondria can also become degraded in vacuoles, and are transported to the organelle through autophagy, as shown by Li et al. (2014). Nonetheless, concentric membranes have apparently not been shown in mitochondria, or in autophagous structures that engulf mitochondria.

In *Dendrobium* tepal cells, mitochondria slowly became devoid of inner membranes, increased in size, and ended as a vacuole-like organelle. Mitochondria at late stages of development apparently did not show an increase in internal membrane profiles (Kirasak et al., 2010). These data mean that it is unlikely that MLBs derive from mitochondria.

### Other possible sources of MLBs

Peroxisomes apparently have not been found to contain concentric membrane structures similar to MLBs, and are therefore likely not precursors. MLBs also are probably not produced by ER membranes as no concentric arrangements of ER have apparently been reported inside the cytosol, and autophagous structures engulfing ER also seems not reported to date. Golgi bodies contain membranes that are often curved, but again no intermediate stages have apparently been reported of Golgi and MLB. This suggests that that MLBs probably do not derive from ER or Golgi bodies.

## Conclusions

The data suggest that MLBs in plants serve to transport membranes to vacuoles, where they will be degraded. This is quite different from the function in animals which often entails deposition to the cell exterior.

The origin of plant MLBs is as yet unknown. The present data seem not to exclude any possible origin. It is here suggested that none of the present data favour the idea that MLBs derive from the ER, from Golgi bodies, or from mitochondria. There is preliminary evidence, it seems, suggesting that MLBs are produced by autophagy, but the role of autophagy might also be limited to the final stage of MLB degradation, whereby fully developed MLBs are taken up by an autophagous structure. Other data suggest the hypothesis that MLBs derive from plastids. The main argument is that structures resembling MLBs are often observed in degenerating chloroplasts and other plastids.

## Acknowledgements

We thank Gabriele Tani and Pietro Di Falco for their fundamental laboratory support. Research funded by the Italian Ministry of the University (MIUR-Fondi di Ateneo)

